# Extracellular matrix sensing via modulation of orientational order of integrins and F-actin in focal adhesions

**DOI:** 10.1101/2022.07.22.501116

**Authors:** Valeriia Grudtsyna, Swathi Packirisamy, Tamara Bidone, Vinay Swaminathan

## Abstract

Specificity of cellular responses to distinct cues from the extracellular matrix (ECM) requires precise and sensitive decoding of information from the cell surface. However, how known mechanisms of mechanosensing such as force dependent catch bonds and conformational changes in focal adhesion (FA) proteins can confer this sensitivity is not known. Using a combination of polarization microscopy and computational modeling, here we identify regulation of orientational order or molecular co-alignment of FA proteins as a mechanism able to precisely tune cell sensitivity to the ECM. We find that αV integrins and F-actin in FAs show changes in orientational order in an ECM-mediated integrin activation dependent manner. This magnitude of orientational order is sensitive to changes in ECM density but independent of myosin-II activity though actomyosin contractility can further fine-tune it. A molecular clutch model for integrin binding ECM ligands demonstrates that orientational order of integrin-ECM binding and catch bonds tune cellular sensitivity to ECM density. This mechanism is also able to decouple ECM density changes from changes in ECM stiffness thus also contributing to specificity. Taken together, our results suggest relative geometric organization of FA components as an important regulator of mechanotransduction.

## Introduction

Cells sense and respond to a wide range of physical cues from the extracellular matrix (ECM) to regulate processes such as cell migration, proliferation, and transcription[1]–[3]. These cues which include the composition and density of ECM proteins and associated mechanical properties such as stiffness, viscoelasticity and architecture are converted into physical signals such as forces and deformation at focal adhesion (FA) sites which are then transmitted and transduced to intracellular protein networks and downstream pathways via the process of mechanotransduction[4]–[6]. Forces and deformation as signals from FAs comprise information about magnitude, direction as well as rates and frequency [7]–[11]. To decipher these signals with high sensitivity and precision, cells need to be able to sense and transmit small changes in the magnitude, direction or loading rates along mechanotransduction pathways while maintaining fidelity of the signal intracellularly.

Over the years, a number of mechanotransduction mechanisms have been discovered, many of which rely on proteins undergoing force-induced changes in their function[12]–[14]. These mechanosensory proteins can modulate their function by exhibiting force-dependent catch bond behavior with interacting partners or by undergoing conformational changes due to application of forces which results in altered protein-protein interactions, modifications or activation[15]–[20]. However, how these mechanisms which primarily rely on forces of different magnitudes acting across the molecule, confer specificity to different physical signals from the ECM is not known. In addition, how sensitivity to different properties of physical signals, such as direction and loading rate is exactly sensed and transduced remains largely unknown.

Recent studies have shown that physical coupling between these different mechanisms at FA sites can indeed bestow sensitivity to changes in ECM stiffness [21]–[23]. Additionally, some catch-bonds in FAs are sensitive to the direction of force acting across the interaction surface of the binding partners[24], [25]. These results imply that instead of acting in isolation, mechanosensors form physical circuits along the force transmission pathway and the relative organization of these molecules with respect to each other as well as with respect to the direction of force can result in changes in sensitivity to ECM cues and downstream cellular response.

We previously showed that the primary ECM receptors, integrins can anisotropically organize and co-align with each other in FAs of cells upon activation by its ECM ligand[26]. This organization which resembles orientational order of nematic materials is oriented along the long axis of FA which is also the direction of the force vector at the cell-ECM interface[27], [28]. Additionally, we observed orientational order of F-actin in FAs correlated with integrin order. Based on the above, we hypothesized that the relative organization of FA proteins as well as their degree of alignment relative to each other and the force direction i.e. the orientational order can underlie ECM sensitivity to different physical cues and downstream cellular response.

### ECM ligand density tunes cell spread area and YAP localization in MEFs in an integrin-dependent manner

We first set out to establish the relationship between physical cues from the ECM and cellular response in our mouse embryonic fibroblasts (MEFs) cell line. We focused on ECM density which allows for precise control of integrin activation, is physiologically relevant and is conducive to high-resolution total internal reflection fluorescence microscopy (TIRF)[29]– [32]. We plated MEFs for 4 hours on a series of glass bottom dishes adsorbed with varying concentrations of fibronectin (0.1 ug/ml, 1ug/ml, 10ug/ml) and poly-l-lysine (PLL)(Fig S1a,b). Fibronectin (FN) binds and activates the β1 and β3 integrin receptors while PLL binds to cells in an integrin activation independent manner. To measure local as well as distal to FA cellular response, we focused on the cell spread area, focal adhesion morphology and nuclear localization of the mechanosensitive transcription factor YAP/TAZ. To directly compare ECM density sensing with the better characterized stiffness sensing, we also plated MEFs on polyacrylamide (PA) gels covalently coupled to fibronectin (10 ug/ml) with Youngs moduli of 0.4, 6 and 60 kPa as previously described[23], [33].

Immunofluorescence staining showed that overall cell morphology on different FN concentrations on glass were clearly distinct, with cells spreading poorly with many small, dispersed focal adhesions and diffused actin and YAP staining on the lowest fibronectin concentration and PLL similar to the 0.4KPa stiffness PA gels (Figure 1a, Figure S1c). Increasing ligand density had a similar effect as increasing stiffness, with cells spreading more, showing distinct actin stress fibers and focal adhesions and clearer nuclear YAP signal (Figure 1a, S1c).

**Figure 1.**
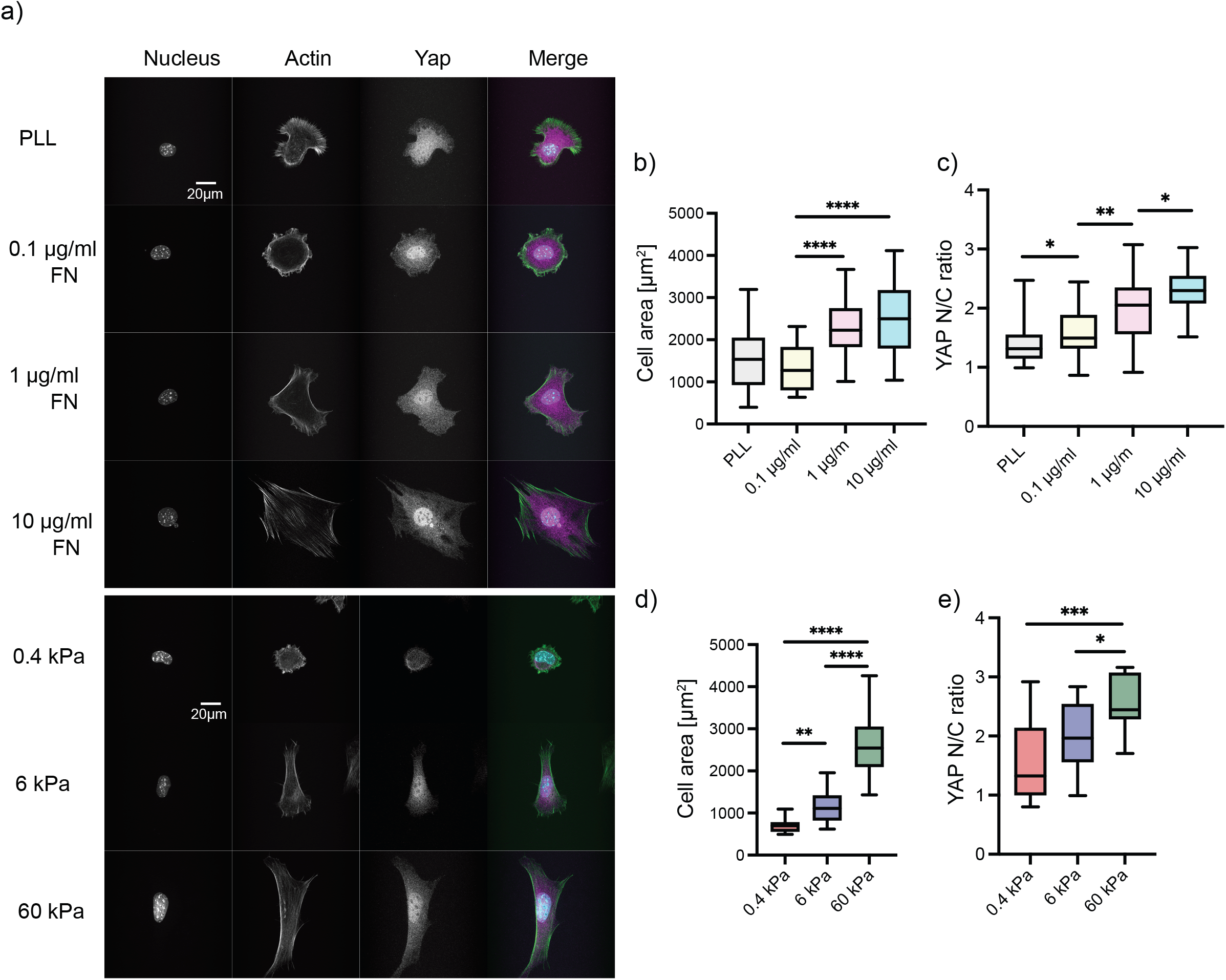
ECM ligand density tunes cell spread area and YAP localization in MEFs in an lntegrin-dependent manner: (a, top) Representative images of MEFs on glass coated with different FN concentrations (0.1 ug/ml, 1ug/ml, 10ug/ml) and PLL, fixed and stained with Hoechst to label nucleus, phalloidin 488 to label actin and Alexa 568 to label YAP. (a, bottom) Representative images of MEFs on gels of different Youngs modulus (0.4, 6, and 60kPa) coated with 10ug/ml of FN, fixed and stained with Hoechst to label nucleus, phalloidin 488 to label actin and Alexa 568 to label YAP. (b) Box plot quantification of cell area from analysis of immunofluorescence images of cells plated on glass coated with different FN concentrations (0.1 ug/ml, 1ug/ml, 10ug/ml) and PLL. Cell area was obtained by segmenting the actin channel. N = 36 for each condition. (c) Box plot quantification of nuclear/cytoplasmic ratio of YAP from analysis of immunofluorescence images of cells plated on glass coated with different FN concentrations (0.1 ug/ml, 1ug/ml, 10ug/ml). N = 36 for each condition. (d) Box plot quantification of cell area from analysis of immunofluorescence images of cells plated on gels of varying Youngs modulus (0.4, 6, and 60kPa), coated with 10ug/ml of FN. Cell area was obtained by segmenting the actin channel. N = 12 for each condition (e) Box plot quantification of nuclear/cytoplasmic ratio of YAP from analysis of immunofluorescence images of cells plated on gels of varying Youngs modulus (0.4, 6, and 60kPa), coated with 10ug/ml of FN. N = 12 for each condition

Quantification of cell spread area based on the actin staining showed a 2 fold increase per 100-fold increase in ligand density while the nuclear to cytoplasmic YAP ratio showed a 1.54 fold increase over these ligand density range (Figure 1b 1c). Quantification of these parameters on PA gels showed a slightly higher level of sensitivity to ECM stiffness with cell area increases by 3.63 from 0.4 to 60KPa and the YAP ratio increasing by 1.85(Figure 1d, 1e). Interestingly, quantification of focal adhesion morphology based on paxillin immunostaining showed small differences in FA size and eccentricity over the range of ligand densities with only the number of focal adhesions/cells being significant (Figure S1, S1d. S1e). We verified the generality of these cellular response in a human epithelial cell line MCF10A as well (Figure S2).

These results collectively show that similar to response to changes to ECM stiffness, cells also respond with high sensitivity to changes in ECM densities in an integrin dependent manner.

### Cellular response to changes in ECM density correlates with changes in integrin activation dependent orientational order of αV integrins and f-actin in FAs

Having previously identified a strong correlation between orientational order of F-actin and αV integrins[26], we first asked if F-actin order was sensitive to changes in integrin activation on FN coated surface. To test this, we plated MEFs untreated or pre-treated with 1 mM Mn^2+^, a potent enhancer of integrin activation, on 0.1ug/ml FN for 4 hours. Cells were then fixed and stained with Alexa Fluor 488-phalloidin and paxillin and imaged using the EA-TIRF system as previously described[26]. Briefly, a polarized excitation is used to illuminate fluorescent molecules in the sample and the parallel (I_‖_) and perpendicular(I_⊥_) components of the emission are separated using a polarized beam splitter. The relative intensities of these 2 components are then used to calculate fluorescence anisotropy using the formula, r= (I_‖_−I_⊥_)/ (I_‖_+ 2I_⊥_). For calculating orientational order of the fluorescent molecule in FAs, the average ‘*r*’ in an FA is plotted as a function of the orientation of the FA long axis with respect to the excitation polarization, across many FAs and cells for a given condition. The amplitude(A) of the modulation in this relationship is dependent on the fraction of molecules co-aligned and directly relates to the molecular order parameter determined for actin in drosophila embryos and the cell membrane[34], [35]. The peak of the modulation in this relationship gives the average orientation of the dipoles of the fluorescent molecules.

EA-TIRFM imaging of the cells showed distinct differences between cells pretreated with Mn^2+^ compared to untreated cells. Mn^2+^ pretreatment increased cell spread area and resulted in more distinct FAs and actin stress fibers compared to the untreated cells (Figure 2a, Figure S2c, S2d)). The paxillin channel was used to segment FAs and the average anisotropy ‘r’ of actin was then calculated per FA and plotted as a function of the FA orientation. Fitting this modulation to a cos^2^ function and extracting the amplitude of modulation showed that indeed integrin activation with Mn^2+^ increased actin order by a factor of 1.78 (A=0.119, R^2^=0.851; A=0.212, R^2^=0.959) (Figure 2b, Figure S3b). Thus, integrin activation increases the orientational order of actin in FAs and confirms that actin order is indeed a consequence as well as a sensitive readout of integrin activation in FAs.

**Figure 2.**
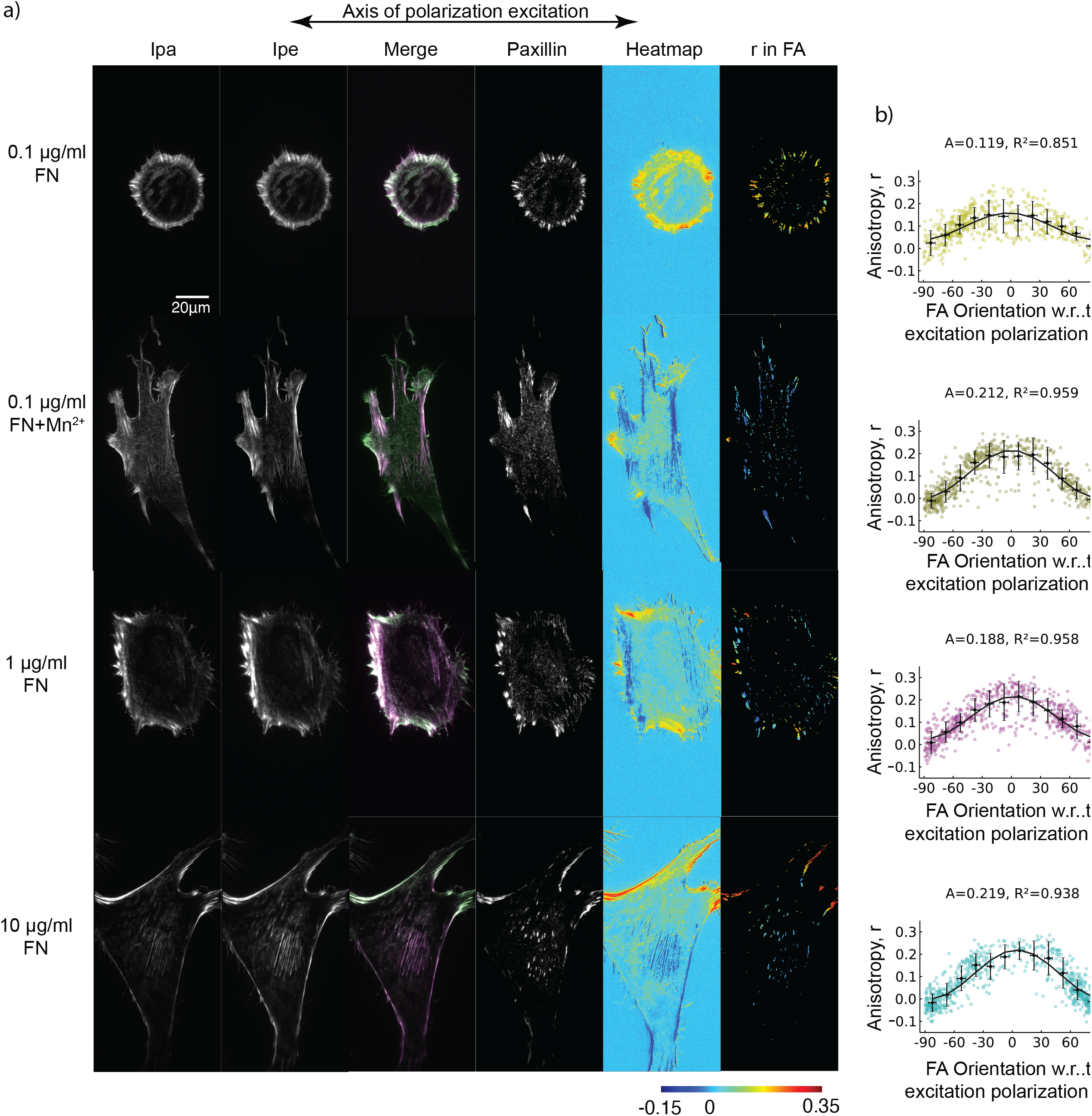
Cellular response to changes in ECM ligand density correlates with finely tuned changes in integrin activation and co-alignment of actin in FAs: (a) Representative images of MEFs on glass coated with different FN concentrations (0.1 ug/ml, 1ug/ml, 10ug/ml) and on 0.1 ug/ml FN in the presence of Mn^2+^. Cells were fixed and stained with phalloidin 488 and imaged with TIRF. Emission from the parallel (Ipa) channel, perpendicular (Ipe) channel and a merge (left, Ipa green, Ipe magenta) are shown. Paxillin stained with Alexa 568 (middle). Emission anisotropy (r) of actin in the whole cell and emission anisotropy of actin in segmented FAs (Right). Magnitude of anisotropy color scale (bottom) (b) mean actin anisotropy (r) in FAs vs FA orientation fit to the cos^2^ function r = C + A·cos^2^(y + θ_d_) for cells plated on each FN condition. Error bars represent SD. Adhesions from 12 cells were analysed.

Next, we asked if ECM ligand density resulted in changes in αV integrin and F-actin order. To measure integrin anisotropy, we expressed a constrained-GFP αV integrin probe which rotationally constrains the GFP to allow measurement of integrin order (Figure 2b,S3b,S4a, S4b). The cells on lower FN concentrations showed similar levels of actin and integrin anisotropy with a narrow dynamic range in all the FAs while cells on higher FN concentration had a higher dynamic range of integrin and actin anisotropy (Figure 2b, Figure S3, S4). The fit of the actin anisotropy vs FA orientation data of cells on 10 ug/ml FN showed a 1.84-fold change in amplitude compared to 0.1ug/ml FN and 1.16-fold change on 1ug/ml FN (Figure 2b, S3b), while the amplitude of integrins across this ECM range changed by a factor of 2.30(Figure S4c). It is important to note that the order parameter for integrins were substantially lower compared to F-actin at the same FN densities.

Thus, taken together, changes in ECM density results in highly sensitive and robust changes in αV integrins and F-actin orientational order which is dependent on integrin activation.

### Myosin II activity is not required but increases F-actin order and enhances cellular sensitivity to changes in ECM density

Within a cell, FAs are highly heterogenous in their size, composition, and function[36]. One of the key regulators of this heterogeneity is the physical coupling between FAs and the actomyosin network which results in differences in tension acting across a FA[37], [38]. During its lifetime, FAs undergo myosin II mediated changes in protein composition, size, and protein organization, all of which are critical determinants of its function[37], [39]–[41]. This suggests that myosin II mediated contractility is a key regulator of FA function. However, it’s role in FA-mediated ECM sensing is controversial as studies have shown that ECM sensing is both dependent [27], [39], [40] and independent [42] of myosin II activity. Here, we next wanted to investigate if orientational order and ECM density sensing was mediated by a specific subpopulation of FAs and thus elucidate the role of myosin II in this organization.

As mentioned above, FA size is strongly correlated with myosin II activity with small adhesions decoupled to contractility and larger FAs coupled to it. So, we re-analyzed our F-actin order measurements under different conditions of FN density by binning all the FAs based on 3 sizes (0.1-0.25μm^2^; 0.25-1μm^2^ and 1-20μm^2^) (Figure 3a). Sizes smaller than 0.1μm^2^ precluded accurate orientation assignment and were thus not analyzed. We then plotted the average ‘r’ as a function of FA orientation for the 3 different size bins and fit the modulations to extract F-actin order at each FN density (Figure 3a). Unsurprisingly, this analysis first revealed that the big FAs (1-20μm^2^) showed significant differences in F-actin order between the lowest FN density (A=0.154; R^2^=0.768) compared to medium and highest FN density (A=0.25; R^2^=0.962) (Figure 3b). However, to our surprise, we also found that the biggest difference in F-actin order at the different ECM densities was in the smallest adhesions (0.1-0.25 μm^2^) with an almost 3-fold change between 0.1ug/ml FN and 1 ug/ml FN and a 5-fold difference between 0.1 to 10 ug/ml FN(Figure 3b). In fact, the cos^2^ fit on the lowest FN density was very poor (R^2^=0.268) leading to an unreliable measure of amplitude and thus suggesting a very disordered actin structure (Figure 3a). Additionally, at the highest FN density, the difference in order of F-actin in the smallest vs. the biggest FAs was very small (1.28-fold change; A=0.196; A=0.25). The middle size bin showed intermediate differences between the small and big FAs with less differences between 0.1 ug/ml FN and 1 ug/ml FN while a 2-fold change between the small and the biggest FAs. Taken together, these results show that that the smallest FAs are the most sensitive to changes in ECM ligand density and that on the highest density, F-actin exhibits orientational order to a similar extent at all FA sizes.

**Figure 3.**
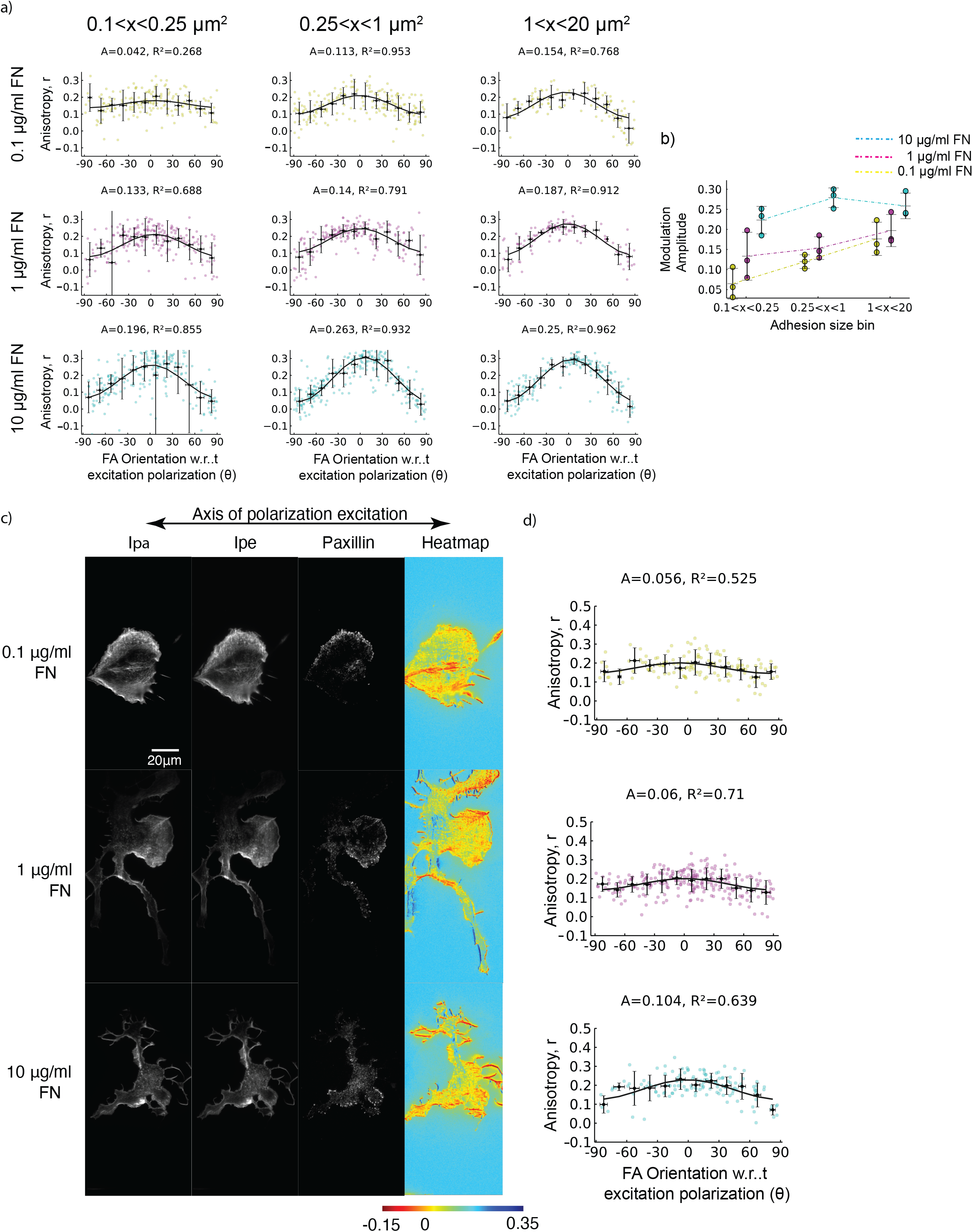
Myosin-2 activity is not required for but enhances ECM density dependent F-actin co-alignment and cellular response: (a) mean actin anisotropy (r) in FAs vs FA orientation fit to the cos^2^ function r = C + A·cos^2^(γ + θ_d_) binned by FA size (0.1-0.25μm^2^; 0.25-1μm^2^ and 1-20μm^2^) in cells plated on glass coated with different FN concentrations (0.1 ug/ml, 1ug/ml, 10ug/ml). (b) Graph showing amplitudes of actin anisotropy (r) binned by FA size from three experiments. (c) Representative images of MEFs on glass coated with different FN concentrations (0.1 ug/ml, 1ug/ml, 10ug/ml) following blebbistatin washout for 5 minutes. Cells were fixed and stained with phalloidin 488 and imaged with TIRF. Emission from the parallel (Ipa) channel, perpendicular (Ipe) channel are shown (left). Paxillin stained with Alexa 568 (middle right). Emission anisotropy (r) of actin stained with phalloidin 488 in the whole cell (right). Magnitude of anisotropy color scale (bottom) (d) mean actin anisotropy (r) in FAs vs FA orientation fit to the cos^2^ function for cells plated on glass coated with different FN concentrations (0.1 ug/ml, 1ug/ml, 10ug/ml) with 5 minutes blebbistatin washout. Adhesions from 15, 12 and 10 cells were analysed for the conditions 0.1 ug/ml, 1ug/ml, 10ug/ml respectively.

Since FA size correlates with FA maturation state and myosin II activity, our results suggested that myosin II activity was not required for changes in ECM density dependent F-actin order. To test this, we plated MEFs on the 3 FN densities and treated cells with the myosin II inhibitor blebbistatin. Inhibition of myosin II results in loss of all stress fibers and big FAs and leaves behind small NAs. However, since NAs are often harder to accurately segment and assign orientation, we washed out the blebbistatin for 5 minutes and then fixed and stained the cells with AF-488 phalloidin and paxillin to analyze F-actin order. EA-TIRFM of actin and analysis on the remaining small FAs after blebbistatin washout revealed that myosin II inhibition resulted in lower F-actin order at all FN densities compared to the untreated cells from Fig 2 (Figure 3c,3d). However, loss of myosin II activity did not completely abrogate FN density dependent changes in order as there still was a 2-fold increase in F-actin order between the 0.1 ug/ml and 10ug/ml (Figure S3a). Thus, ECM density dependent F-actin order is independent of myosin II activity, though its activity can increase this order.

The above results suggest that cellular response to changes in ECM ligand density is independent of myosin II activity with potentially a role for myosin II in enhancing sensitivity to these changes. To test this, we plated MEFs on FN coated cover-glass at the 3 densities and then pretreated the cells with blebbistatin or with DMSO (Figure 4a). We fixed and stained the cells for either actin to measure cell spread area or with YAP to measure its nuclear translocation. Quantification and analysis showed an insignificant change in cell spread area in blebbistatin treated cells compared to the DMSO control over each concentration of FN (Figure 4b). This led to overall similar sensitivity in cell area across the different ECM ligand densities. Myosin II inhibition also led to decrease in YAP nuclear translocation at 0.1ug/ml FN and 10 ug/ml FN, while there was no statistical change at 1 ug/ml FN (Figure 4c). This led to a slight decrease in ECM ligand sensitivity in overall YAP nuclear translocation with a 1.26-fold change from the lowest to the highest(Figure 4c).

**Figure 4.**
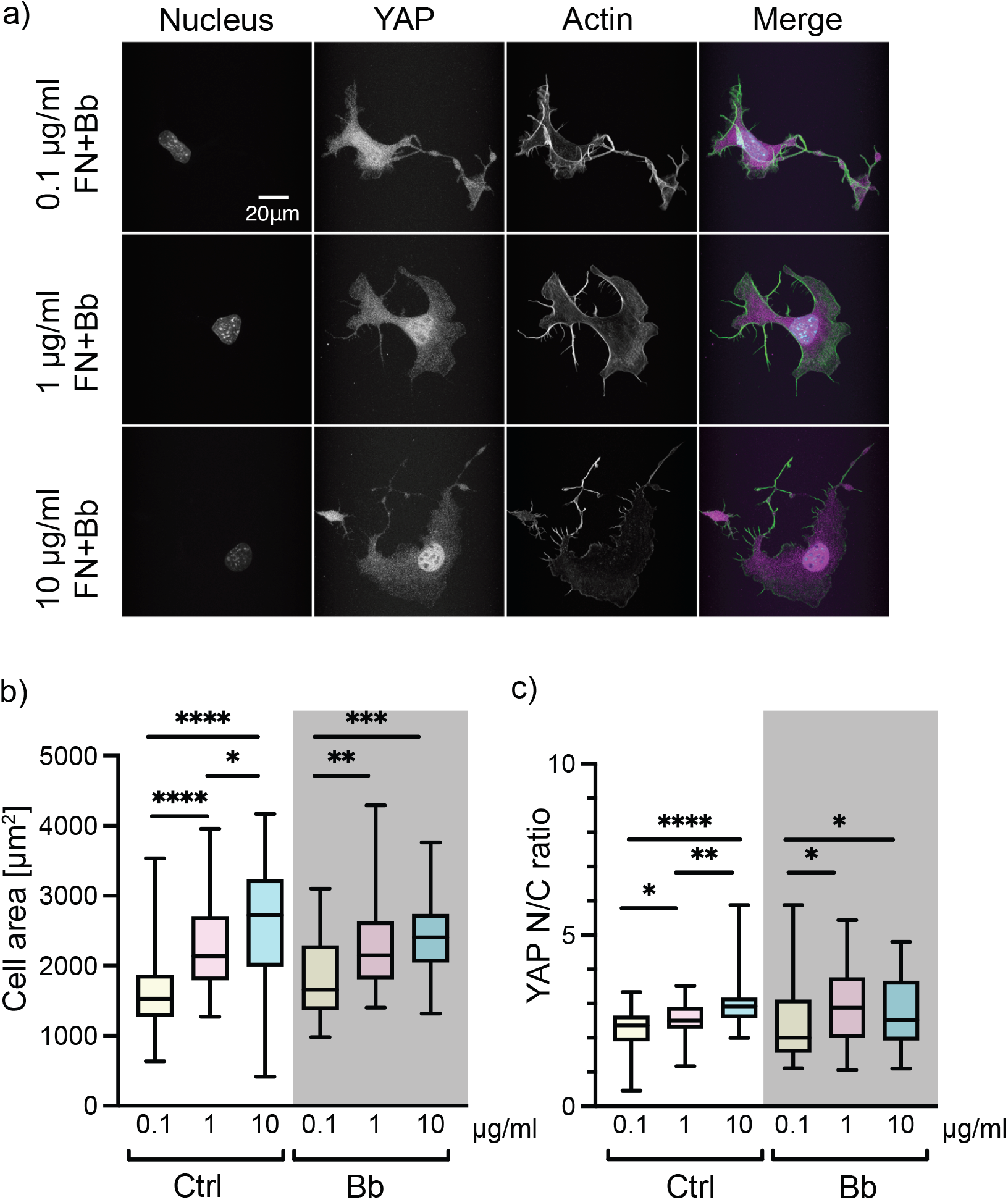
F-actin anisotropy in FAs is myosin 2 independent (a) Representative images of MEFs on glass coated with different FN concentrations (0.1 ug/ml, 1ug/ml, 10ug/ml) following blebbistatin treatment, fixed and stained with Hoechst to label nucleus, phalloidin 488 to label actin and Alexa 568 to label YAP. (b) Box plot quantification of cell area from analysis of immunofluorescence images of cells plated on glass coated with different FN concentrations (0.1 ug/ml, 1ug/ml, 10ug/ml) following blebbistatin treatment. Cell area was obtained by segmenting the actin channel. N = 36 for each condition. (c) Box plot quantification of nuclear/cytoplasmic ratio of YAP from analysis of immunofluorescence images of cells plated on glass coated with different FN concentrations (0.1 ug/ml, 1ug/ml, 10ug/ml) following blebbistatin treatment. N = 36 for each condition

Taken together, these results suggest that ECM ligand density dependent changes in orientational order of F-actin and cellular response is independent of myosin II activity, though motor activity can increase orientational order and enhance cellular sensitivity.

### Orientational order of FA components increases sensitivity for ECM ligand binding by modulating directional catch bonds

We next evaluated what mechanisms underlie the changes in orientational order of F-actin to enable sensitivity to ECM density independent of myosin II activity. To do so, we extended our computational model of nascent adhesion assembly based on the molecular-clutch mechanism [43], [44]. Each clutch in the model was represented as an explicit particle diffusing on the top surface of two parallel surfaces, with the top surface mimicking the ventral surface of a cell, and the bottom one mimicking the substrate with randomly distributed ligands (Figure 5a). Each clutch underwent cycles of diffusion, binding and unbinding of substrate ligands.

**Figure 5.**
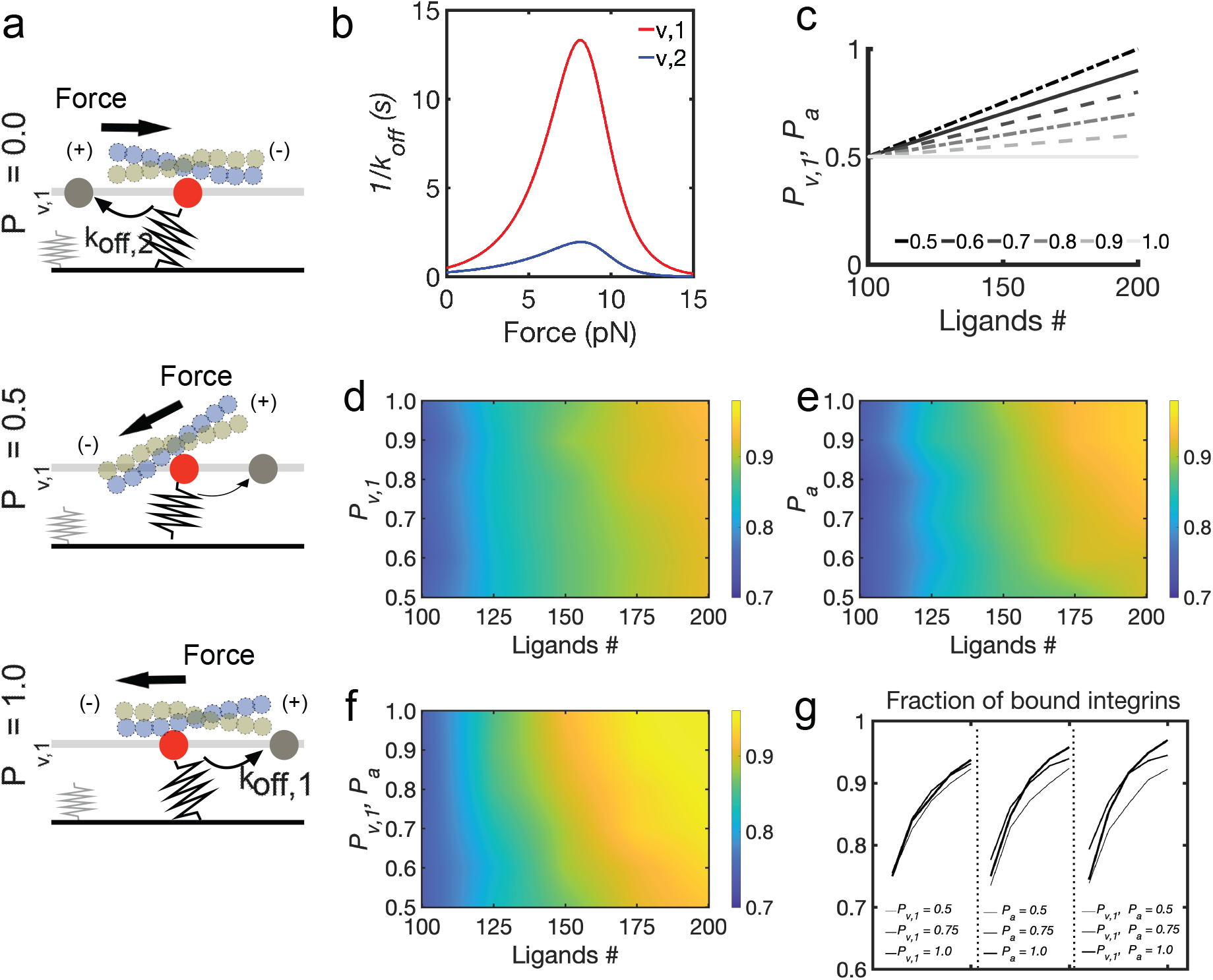
Ligand sensitivity is governed by molecular order of FA clutches. (a) Schematic representation of the computational model of FA assembly based on the molecular clutch mechanism. Two 2D surfaces are placed 20 nm apart. The bottom surface represents the substrate with a random distribution of ligands, modeled as elastic springs with stiffness *k*_*sub*_, which depends on the substrate young modulus, *Y*. The top surface mimics the ventral cell membrane, with integrins diffusing with diffusion coefficient *D* and establishing interactions with substrate ligands. When integrin binds a substrate ligand, filaments exert a force on the clutch with magnitude that depends on their retrograde velocity and myosin motors activity. The direction of this force determines the clutch unbinding rate, *k*_*off,1*_ (directional) or *k*_*off,2*_ (non-directional). (b) Unbinding rates follow the lifetime (1/*k*_*off*_) versus force relation, where the directional pathway of vinculins in the clutch is indicated with v,1, and the non-directional pathway is indicated with v,2. (c) Linear relations between the probability of *k*_*off,1*_ (directional pathway for unbinding) *P*_*v,1*_, and integrin activation rate, *P*_*a*_, with varying ligand density, *n*, between 100-200 ligands/um^2^. Legends indicate the value of *Psub>v*,_*1*_ or *P*_*a*_ using *n* = 200. The following relations are used: for *P*_*v,1*_ or *P*_*a*_ = 0.5 at *n* = 200, P_v,1_ or *P*_*a*_ = 0.0 x *n* +0.5; for *P*_*v,1*_ or *P*_*a*_ = 0.6 at *n* = 200, *P*_*v,1*_ or *P*_*a*_ = 0.01 x *n* +0.4; for P_v,1_ or *P*_*a*_ = 0.7 at *n* = 200, *P*_*v,1*_ or *P*_*a*_ = 0.02 x *n* +0.3; for *P*_*v,1*_ or *P*_*a*_ = 0.8 at *n* = 200, P_v,1_ or *P*_*a*_ = 0.03 x *n* +0.2; for *P*_*v,1*_ or *P*_*a*_ = 0.9 at *n* = 200, *P*_*v,1*_ or *P*_*a*_ = 0.04 x *n* +0.1; for P_v,1_ or *P*_*a*_ = 1 at *n* = 200, *P*_*v,1*_ or *P*_*a*_ = 0.05 x *n*; (d) Average fraction of ligated clutches varying *P*_*v,1*_ between 0.5-1 and n between 100-200 ligands/um^2^, while keeping *Pa* = 0.5. (e) Average fraction of ligated clutches varying *P a* between 0.5-1 and n between 100-200 ligands/um^2^, while keeping *P*_*v,1*_ = 0.5. (f) Average fraction of ligated clutches by simultaneously varying *P*_*v,1*_ and *P*_*a*_ between 0.5-1 and varying *n* between 100-200 ligands/um^2^. (g) Average fraction of ligand-bound integrins for different values of *P*_*v,1*_, *P*_*a*_ or both, varying *n* between 100-200 ligands/um^2^. All data are obtained as averages from 300 s of simulations for each condition, using *Y* = 6 KPa.

Since our experiments showed that αV integrins orientational order increases with increase in ECM density (Figure S3), we incorporated a probability of integrin activation (*P*_*a*_) that varies linearly with ECM density, with a minimum of 0.5 at 100 ligands/um^2^ and a maximum of 1 at 200 ligands/um^2^ (Figure 5c). When integrin was active, the clutch could bind a free ligand; when integrin was inactive it could only diffuse. When bound to a ligand, each clutch was also considered bound to the actin cytoskeleton through binding of vinculin to actin (via talin). While the mechanism by which orientation order of F-actin is established in FAs is unknown, a previous study has shown that long range F-actin order can be established by formation of directional catch bonds between vinculin and F-actin [25]. Having explicitly measured orientational order of F-actin here (Figure 2), we incorporated in the model changes in F-actin order by using two pathways for clutch unbinding of actin: directional, *k*_*off,1*_, and non-directional, *k*_*off,2*_ (Figure 5a-b). The directional pathway corresponded to the one with the highest peak in bond lifetime for the vinculin-actin bond at ∼13 s (Figure 5b), in which vinculin forms a bond with actin pulling towards its pointed end. The non-directional pathway corresponded to the lifetime-force relation with a maximum of ∼ 3s (Figure 5b). To include changes in orientational order of F-actin, we tuned the probability of clutch unbinding actin according to the directional unbinding pathway (*P*_*v,1*_) and then tested its importance in mediating ligand sensitivity (Figure 5c). To evaluate how different ECM-dependent integrin anisotropy could affect adhesion assembly, we tuned the degree by which *P*_*a*_ varies with ligands concentration (Figure 5d) and evaluated its effect on the fraction of ligated clutches. Last, to understand how the combined effects providing orientational order of FA clutches, ECM-dependent integrin activation and F-actin orientation, affect sensing of ECM density, we tested how simultaneously varying *P*_*v,1*_, and *P*_*a*_, affected the average number of ligated clutches (Figure 5e). Varying the maximum *P*_*v,1*_ (*P*_*v,1*_ corresponding to *n* = 200 ligands/um^2^) from 0.5 to 1 (Figure 5c) while maintaining *P*_*a*_ = 0.5, increased the average fraction of ligated clutches from a minimum of ∼0.73 to a maximum ∼0.93 using 100 to 200 ligands/um^2^(Figure 5d). Varying the maximum *P*_*a*_ (*P*_*a*_ corresponding to *n* = 200 ligands/um^2^) from 0.5 to 1 (Figure 5c) and varying *n* from 100 to 200 ligands/um^2^, while maintaining *P*_*v,1*_ = 0.5, decreased the minimum fraction of ligated clutches to ∼0.71 and increased their maximum to ∼ 0.95 (Figure 5e).

Simultaneously varying *P*_*a*_ and *P*_*v,1*_ from 0.5-1 (Figure 5c) in the same range of ligand densities resulted in variations of the fraction of ligated clutches from 0.73, up to ∼ 0.97 (Figure 5f). The fraction of ligated clutches increased about 1.6% using *n* = 200 ligands/um^2^ and varying *P*_*v,1*_ from 0.5 to 1, while keeping *P*_*a*_ = 0.5 (Figure 5g). The fraction of ligated clutches increased by about 4% using *n* = 200 ligands/um^2^ and varying *P*_*a*_ from 0.5 to 1, while keeping *P*_*v,1*_ = 0.5 (Figure 5g). By simultaneously increasing both *P*_*a*_ and *P*_*v,1*_ this increase in the fraction of ligated clutches was more than 5%. The percentage of short-lived clutches was lower using *P*_*a*_ = *P*_*v,1*_ = 1 than using either *P*_*a*_ = 0.5 or *P*_*v,1*_ = 0.5 (Figure S6A), meaning that clutches were ligated for longer time when FA order was maximized. Collectively, these results demonstrate that ligand sensitivity increases the most by increasing both in the probabilities of integrin activation and directional catch bond. By contrast, by increasing either parameter in isolation narrows the range of the fraction of ligated clutches, resulting in less sensitivity to ECM density. Therefore, the results from the model demonstrate that the magnitude of orientational order of actin filaments determine the precise sensitivity to ECM density through FA unbinding kinetics in an integrin-activation dependent way.

We next wondered if ECM density sensing was distinct and independent of ECM stiffness in our orientational order-based motor clutch model. To do so, we systematically changed the ECM stiffness from 0.4kPa to 60kPa and plotted the distribution of the fraction of ligated clutches at 3 different ECM ligand densities (100, 110 and 180 ligands/um^2^) (Figure 6a). Using 0.4kPa, the fraction of ligated clutches increased from ∼0.6 to ∼0.8; using 6 kPa the fraction of ligated clutches increased from ∼0.7 to ∼0.95, shifting the range of ligand sensitivity upward; using 60 kPa, the fraction of ligated clutches increased from only ∼0.55 to ∼0.85, shifting the range of ligand sensitivity downward. Surprisingly, ECM density sensing was independent of ECM stiffness in our model as all stiffnesses shown a response to changes in ligand density. At different stiffnesses, sensitivity to ECM density also increased the most when *P*_*a*_ and *P*_*v,1*_ were increased simultaneously (Figure S6). Interestingly, our model also showed a biphasic response across different stiffnesses for the same ligand density (Figure 6a). This occurred because when substrate forces were too high adhesions disassembled instead of stabilizing.

**Figure 6.**
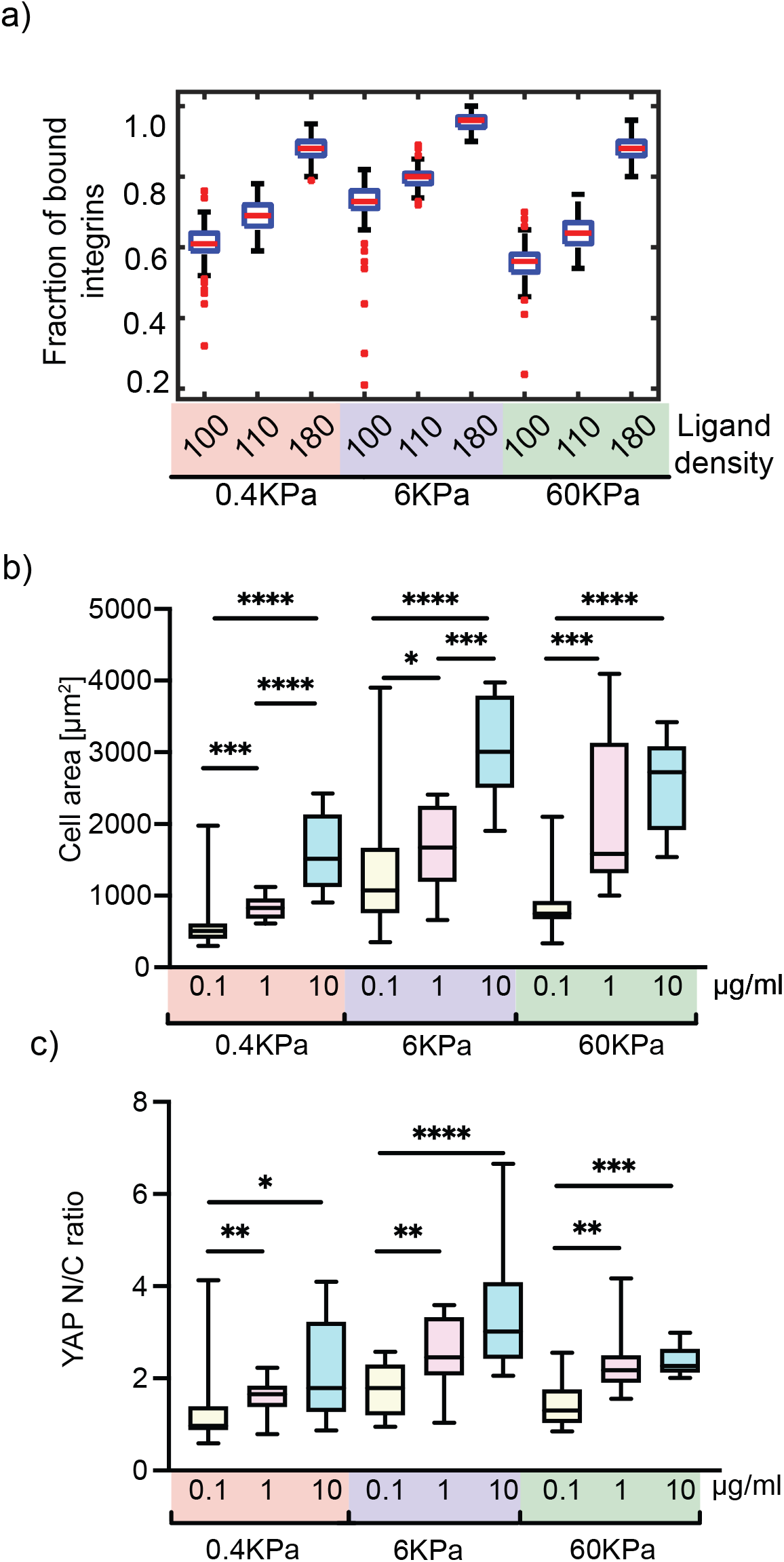
ECM ligand sensitivity is ECM stiffness independent: (a) Boxplots of the fraction of ligand bound integrins varying n between 100-180 ligands/um^2^ and using Y= 0.4, 6, and 60 kPa. All data are obtained from 300 s of simulations sampling every 1 s. (b) Box plot quantification of cell area from analysis of immunofluorescence images of cells plated on gels of varying Youngs modulus (0.4, 6, and 60kPa) coated with different FN concentrations (0.1 ug/ml, 1ug/ml, 10ug/ml). Cell area was obtained by segmenting the actin channel. N = 12 for each condition. (c) Box plot quantification of nuclear/cytoplasmic ratio of YAP from analysis of immunofluorescence images of cells plated on gels of varying Youngs modulus (0.4, 6, and 60kPa) coated with different FN concentrations (0.1 ug/ml, 1ug/ml, 10ug/ml). N = 12 for each condition

We tested these predictions of our model by plating MEFs on the 3 different FN densities (0.1-10 ug/ml) on PA gels of the 3 stiffnesses used in the model (0.4 KPa, 6 KPa and 60 KPa). We fixed and immunostained the cells after 4 hours of plating for YAP, actin and nucleus, just as previously described. Quantification of the cell spread area (Figure 6b) and YAP N/C ratio (Figure 6c) across all these conditions showed a remarkable match with the prediction from the model. At each ECM rigidity, cell area and nuclear localization of YAP increased with increasing FN density with different sensitivities. We also observed a biphasic response at the highest ECM density (10ug/ml) across the 3 rigidities matching the prediction of the model.

Taken together, these results confirm that ECM dependent changes in orientational order of FA components and regulation of directional catch bonds provide a robust mechanism for fine tuning the sensitivity of cells to ECM density, independent of ECM rigidity.

### Outlook

Integrin-based FAs are the primary multimolecular structures that mediate sensing of a wide range of physical cues from the ECM. At the molecular scale, FAs perform this function by assembling a network of several functionally different proteins that are functionally and physically linked[37], [45], [46]. In addition to composition, FA molecular architecture is also highly complex and thought to be a critical regulator of its function[47]–[49]. The role of multimolecular architecture is even more important if one considers the mechanisms of activation of mechanosensory molecules essential for FA function. The changes in protein function of mechanosensors are driven by changes in protein structure or conformation upon application of forces which expose new binding surfaces or cryptic sites that undergo further modifications. Owing to physical properties of force such as magnitude and direction, the geometry or orientation of interactions between forces and proteins is an important factor that can influence the changes in conformation in protein structure and thus its function[50], [51]. In addition to its effect on single proteins or cluster of proteins, the relative orientational organization of different proteins, physically connected across the whole FA or FA-nucleus or FA-other cellular structures can have an impact on the nature of forces felt by distal proteins and organelles[52]. However, the link between physical cues from the ECM, molecular organization within the FA and cellular response has so far been missing.

Here, we have shown that FA molecules can exhibit orientational order which most importantly, changes upon small changes in physical cues from the ECM. Our data on F-actin and integrins in combination with our previously published data shows that FAs have an anisotropic molecular architecture that is dynamic and originates at the ECM-integrin interface and extends to the actin cytoskeleton, which is indirectly connected to integrins. We find that the establishment and regulation of order does not require myosin II contractility suggesting that forces due to actin polymerization is sufficient and thus not required for mechanosensing, though contractility can enhance both order and mechanosensitivity. Currently, the specific mechanism that results in F-actin order in FAs is unknown. A computational model for directionally asymmetric catch bonds between vinculin and F-actin has previously shown the establishment long range F-actin order[25]. In our study, we show that the modulation of orientational order measured here when linked to the directional catch bond as previously shown can lead to highly sensitive mechanosensing of ECM cues. However, it also likely that other mechanisms can contribute to establishment and regulation of orientational order, which include other directional catch bonds as well as actin crosslinking proteins that have shown to promote actin order in focal adhesions[24], [53], [54]. Interestingly, long range order of actin in cells has been previously shown to play a significant role in modulating ECM dependent changes in its mechanical properties[55]. While it’s tempting to speculate that the origin of this long range order is at FAs via the mechanism or orientational order discovered here, this and other regulatory mechanisms will be the subject of future research.

Modulation of orientational order also offers an elegant and energetically less expensive mechanism for enhancing sensitivity to ECM cues compared to recruitment of new proteins to the FAs and is likely applicable to other subcellular structures as well where previously, order of protein components have been observed[56], [57]. In addition, these results further emphasis unique material properties of FAs which seem to resemble liquid-crystalline materials rather than disordered liquid-like. This may have implication in the assembly and growth of these structures in the cell as well as in-vitro reconstitution efforts which utilize phase separation methodologies[58], [59]. Finally, orientational order of load bearing proteins have been observed across different cell types, length scales and the animal kingdom[60]–[62]. In combination with the results of this study, this suggests a potentially conserved mechanism where orientational order is not just the consequence of directional forces but may also be a mechanism that fine tunes response to the forces across different cell and tissues in our body.

## Methods

### Cell culture and sample preparation

Mouse embryonic fibroblasts (MEFs) were cultured in Dulbecco’s high glucose modified eagle medium (Gibco) supplemented with 10% fetal bovine serum (Gibco) and 100 units/ml Penicillin/Streptomycin (Gibco). Polyacrylamide gels of varying Youngs modulus were prepared and functionalized following a previously described protocol [63]. Functionalized polyacrylamide gels or 35 mm 1.5 glass-bottom dishes (Cellvis) were coated with Poly-L-lysine (PLL) (Sigma-Aldrich), 0.1, 1 and 10 ug/ml fibronectin (Sigma-Aldrich) for 30 min at 37°C and blocked with 2% Bovine Serum Albumin (Sigma-Aldrich) in PBS for 1hr at 37°C or at 4°C overnight. Cells were plated on FN/PLL coated glass-bottom dishes or gels for 4h prior to fixation and immunostaining. The relative change in ECM density was verified by staining for fibronectin (Figure S1). For blebbistatin washout experiments, cells were allowed to attach for 2h prior to adding 50μM blebbistatin or DMSO which was washed out after 2h for 5min. For integrin activation experiments, cells were plated in the presence of 1mM Mn^2+^ for 4h before fixing and staining.

### Immunostaining and transfection

Cells were fixed with 4% paraformaldehyde (Thermo Scientific) in cytoskeletal buffer (CB) (2 mM EGTA, 138 mM KCl, 3 mM MgCl, 10 mM Mes, pH 6.1) for 20 min at 37°C and permeabilized with 0.5% Triton TX-100 (Sigma-Aldrich) in CB for 5 min at RT. They were then incubated with 0.1M glycine in CB for 10 min and washed with Tris Buffered Saline buffer (TBS), once for 5 min and twice for 10 min. Next, the cells were blocked with 2% BSA in 0.1% tween TBS (TBST) for 1h and immunostained with mouse anti-paxillin IgG1 antibody (BD Transduction Laboratories 610052) (1:500) overnight at 4°C. The cells were washed with TBST and incubated with secondary goat anti-mouse IgG Alexa 568 (Invitrogen A11031) (1:400) and Alexa Fluor 488 Phalloidin (Invitrogen) (1:500) for 2h. Samples were washed and imaged in TBS or mounted on glass slides with ProLong Glass Antifade Mountant (Invitrogen).

For YAP immunostaining and cell area measurements, cells were fixed with 4% paraformaldehyde in PBS for 15 min at RT, permeabilized with 0.2 % Triton TX-100 in PBS for 5 min and blocked with 3% BSA in PBS for 45 min. Cells were immunostained with primary mouse-anti YAP monoclonal IgG2a antibody (Santa Cruz biotechnology sc-101199) (1:400) in 1% BSA in PBS and incubated at 4°C overnight. Cells were washed with 0.05% Tween in PBS and incubated for 2h in secondary goat anti-mouse Alexa 568-conjugated IgG antibody (Life Technologies Corporation), Alexa Fluor 488 Phalloidin (Invitrogen) (1:500) and nuclear dye Hoechst 350 (33342 Solution, Thermo Scientific) (1:4000). Samples were washed and imaged in TBS or mounted on glass slides.

To measure anisotropy and orientational order of αV Integrin, αV Integrin-GFP-constrained plasmid based on previous work was used[26]. Cells were co-transfected using the Neon Transfection system (Invitrogen MPK5000) with 2500ng of αV Integrin-GFP-constrained plasmid and untagged β3 integrin plasmid at 1500V, 30ms, 1pulse. Transfected cells were cultured in antibiotic free media for 48h before plating on FN/PLL coated 35 mm 1.5 glass-bottom dishes for 4h. The cells were fixed and stained for paxillin.

### EA-TIRFM, image acquisition

Images were acquired using total internal reflection fluorescence microscopy (TIRF) configuration on a Nikon Eclipse Ti microscope with the following available laser lines: 405 nm, 488 nm, 561 nm and 657 nm and Spectra EX (Lumencor). TIRF APO 100x 1.49 N.A. objective was used for acquiring images. Emission/excitation filters used were: GFP (mirror: 498-537 nm and 565-624nm; excitation: 450-490 nm and 545-555 nm; emission: peak 525 nm, range 30 nm) and mCherry(mirror: 430-470 nm, 501-539 nm and 567-627 nm; excitation: 395-415 nm, 475-495 nm and 540-560 nm; emission: peak 605nm, range 15 nm) or Continuous STORM (mirror: 420-481 nm, 497-553 nm, 575-628 nm and 667-792 nm; excitation: 387-417 nm, 483-494 nm, 557-570 nm and 636-661 nm; emission: 422-478 nm, 502-549 nm, 581-625 nm and 674-786nm).

A polarized evanescent TIRF wave excited the fluorophores. In the emission pathway, the emitted light was split into p and s polarization components with a polarization beam splitter (Laser Beamsplitter zt 561 sprdc), placed into an Optosplit III. The orthogonal images (Ipa and Ipe) were projected onto two separate fields of view, manually aligned and captured with a Teledyne Photometrics 95B 22mm camera. The polarization of the evanescent field was verified by measuring the extinction coefficient.

### EA-TIRFM image analysis

All image analysis were performed using Julia (v.1.6.0) and the code is available on request. The Ipa and Ipe images were aligned using QuadDIRECT (https://github.com/timholy/QuadDIRECT.jl). G-factor, 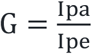 was calculated in conjunction with every imaging session to correct for polarization bias in the detection system and the optical path. A low concentration solution of fluorescein (Sigma-Aldrich) in water was imaged with the same camera settings, using 488 nm LED for excitation [64]. The resulting Ipa and Ipe images were used for G-factor computation.

The fluorescence anisotropy formula,

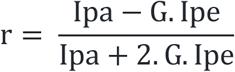

was applied to the orthogonal F-actin-phalloidin or const. αV integrin images to create an anisotropy heatmap. FAs were segmented from the background subtracted paxillin image using the “Moments” binarization algorithm from the ImageBinarization.jl package. The FA mean anisotropy and the angle between the FA long axis and the excitation polarization (θ), were extracted using in-house packages in Julia. FAs with eccentricity less than 0.9 (and less than 0.7 in the Bb washout experiments were filtered out for reliable estimation of FA long axis orientation.

Mean anisotropy values (mean intensities from the anisotropy heatmap) from all FAs, collected from all cell images within the same condition, were plotted against the corresponding θ angles. The data was binned into 15-degree bins (12 bins in total) and the mean ‘r’ together with SD were calculated. The following trigonometric function was fitted to the binned means using the “curve_fit” function from the LsqFit.jl package:

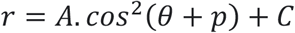

where r is the fluorescence emission anisotropy; amplitude A is directly related to orientational order; θ is the angle between the FA long axis and the excitation polarization; p is the phase shift, which defines the average orientation of dipoles in the ensemble of fluorophores; C is the anisotropy offset due to background. Negative counterclockwise angles were converted to positive angles.

### Confocal microscopy, calculation of YAP N/C ratio and FA analysis

Images were acquired using a Nikon Confocal A1RHD microscope, with 405 nm, 488 nm, 561 nm, and 640 nm laser lines and equipped with 2 GaAsP PMT (for 488 and 561 lines), 2 PMT (for 405 and 640 lines) and 1 GaAsP PMT for spectral detection. Following objectives were used for imaging: TIRF APO 100x 1.49 N.A. and Plan Apochromal Lambda 60x 1.42 N.A. To capture the entire cell volume z-stacks with 0.6 um step size for cells on glass and 1 um step size for cells on gels were taken (usually 10-20 steps per cell).

The maximum projection of the z-stack in YAP channel was background subtracted using functions “sigma_clip” and “estimate_background” provided by the Photometry.jl. Median filtering was applied to images taken with 100x objective. The nucleus was segmented from the summed up nuclear channel z-stack and eroded (with five pixels for the images taken with 60x objective and 7 pixels for images taken with 100x objective) to represent the area with nuclear YAP. The nuclear segment was dilated by 55 (for 60x) or 59 (for 100x) pixels and subtracted with a nuclear segment dilated by 10 (for 60x) or 14 (for 100x) pixels, to attain a ring-shaped segment around the nucleus. This way, the transition area between the nuclear and the cytoplasmic region was excluded. The cell was segmented in the actin channel to represent the cell spread area. The intersection of the ring-shaped segment and slightly eroded actin segment represented the area with cytoplasmic YAP. For computation of N/C YAP ratio the median YAP intensity was divided by the median cytoplasmic YAP intensity. The entire cytoplasmic area was not used to avoid the contribution of low YAP counts caused by thin cell edges. The Julia code is available on request.

Paxillin images, taken as previously described in TIRF mode, were submitted for analysis to the online Focal Adhesion Analysis Server (FAAS)[65].

### Molecular clutch model of adhesion assembly

To understand how the orientational order of actin filaments could enable ligand sensitivity, we extended our computational model of nascent adhesion assembly at the leading cell edge. Like previous approaches from us and others[44], [66]–[68], our model was based on the molecular-clutch mechanisms, in which adhesion clutches intermittently transmit cytoskeletal force to the substrate and substrate rigidity to the actin flow. Cytoskeletal force was produced by intracellular myosin motors regulating the actin flow velocity and the number of engaged clutches. Substrate force regulated the strength of the clutch-ligand binding, mimicking integrin-fibronectin bonds in terms of lifetime versus force relations. Cytoskeletal force regulated the strength of the clutch-actin binding, mimicking vinculin-actin bonds in terms of directional versus non-directional lifetime versus force.

Each clutch in the model was represented as an explicit single point particle in 3D, that included integrin and vinculins. Once a clutch bound a substrate ligand, it was considered also bound to the actin cytoskeleton. The lifetime of the fibronectin and actin bound state of the clutch depended on the kinetics of the bonds of integrin with substrate ligand and of vinculins with actin. It has been shown that both integrin–fibronectin bond and vinculin-actin bond are catch-slip bonds, meaning that their lifetime first increases as a function of force, then decreases as the force increases further[16], [25].

Initially, a given number of fibronectin molecules were randomly distributed on a substrate, to which integrins could bind. The substrate was represented as an isotropic and elastic material, consisting of a bundle of ideal springs which mimicked ligands.

The clutches underwent cycles of diffusion along the top surface of the model mimicking the ventral membrane of cells, followed by binding, and unbinding of substrate ligands underneath it (Figure 5a). As clutches bound substrate ligands, they experienced cytoskeletal force, from vinculins binding actin undergoing retrograde flow, as was seen in the lamellipodium[69]. The number of vinculin-actin bonds depended on force and varied between 2 and 11[70]. When a diffusing clutch came in proximity of a free fibronectin, it established a harmonic interaction, which mimicked binding, with stiffness determined by substrate rigidity. When either integrin or at least one vinculin were bound, the clutch was considered engaged. When the clutch was engaged, it transmitted the actin flow to the substrate and built tension on the bonds. This tension was used to determine the unbinding rates of vinculins and integrin. Therefore, these clutches governed local balances between cytoskeletal contractile force and substrate stiffness. When both integrin and all vinculins in a clutch were unbound, the clutch was free to diffuse, mimicking the free diffusion of integrin receptors on the ventral surface of cells[71]. The actin flow was modulated by the number of clutches engaged with the substrate, and the number of molecular motors in the system. The force on the integrin-fibronectin bonds was also directly proportional to substrate stiffness. All parameters in the model were based upon available experimental data and previous modeling approaches[72]. We directly incorporated the lifetime vs. force relationships of integrin–fibronectin and vinculin-actin bonds from atomic force microscopy and optical trap single-molecule experiments[16], [25]. To account for ligand-dependent integrin activation we varied the probability of integrin activation, *P*_*a*_. To account for different degrees of actin filaments orientation we systematically varied the probabilities of directional catch bond for vinculin, *P*_*v1*_. (Figure 5c). To quantify the amount ligated clutches, we measured the average fraction of ligated clutches from simulations of 300s.

## Data and code availability

All raw data that support the findings and the image analysis codes are available from the corresponding authors upon request.

## Acknowledgements

We thank P Nordenfelt, Amin Doostmohammadi, Martijn Gloerich, Guillaume Jacquemet, Sebastian Wasserstrom and all the members of laboratory of cell and molecular mechanobiology (LCMM) for their discussion and support. Johannes Kumra Ahnlide is specially acknowledged for developing the initial analysis pipeline and all the help in developing code and maintaining image storage servers. Lund University Bioimaging Centre (LBIC) at Lund University is gratefully acknowledged for providing experimental resources. This research was funded by the Knut and Alice Wallenberg foundation (VS, Wallenberg Centre for Molecular Medicine, Lund);Cancerfonden (VS, 19 0445 Pj, Projekt grant) and the National Science Foundation (TCB, NSF BMMB 2044394).

## Author contributions

VG and VSS conceived the project. VG and SP performed experiments and analysed data. TCB developed the computational model. VG, SP, TCB and VS wrote the manuscript. VSS supervised the project.

---

**Figure S1.**
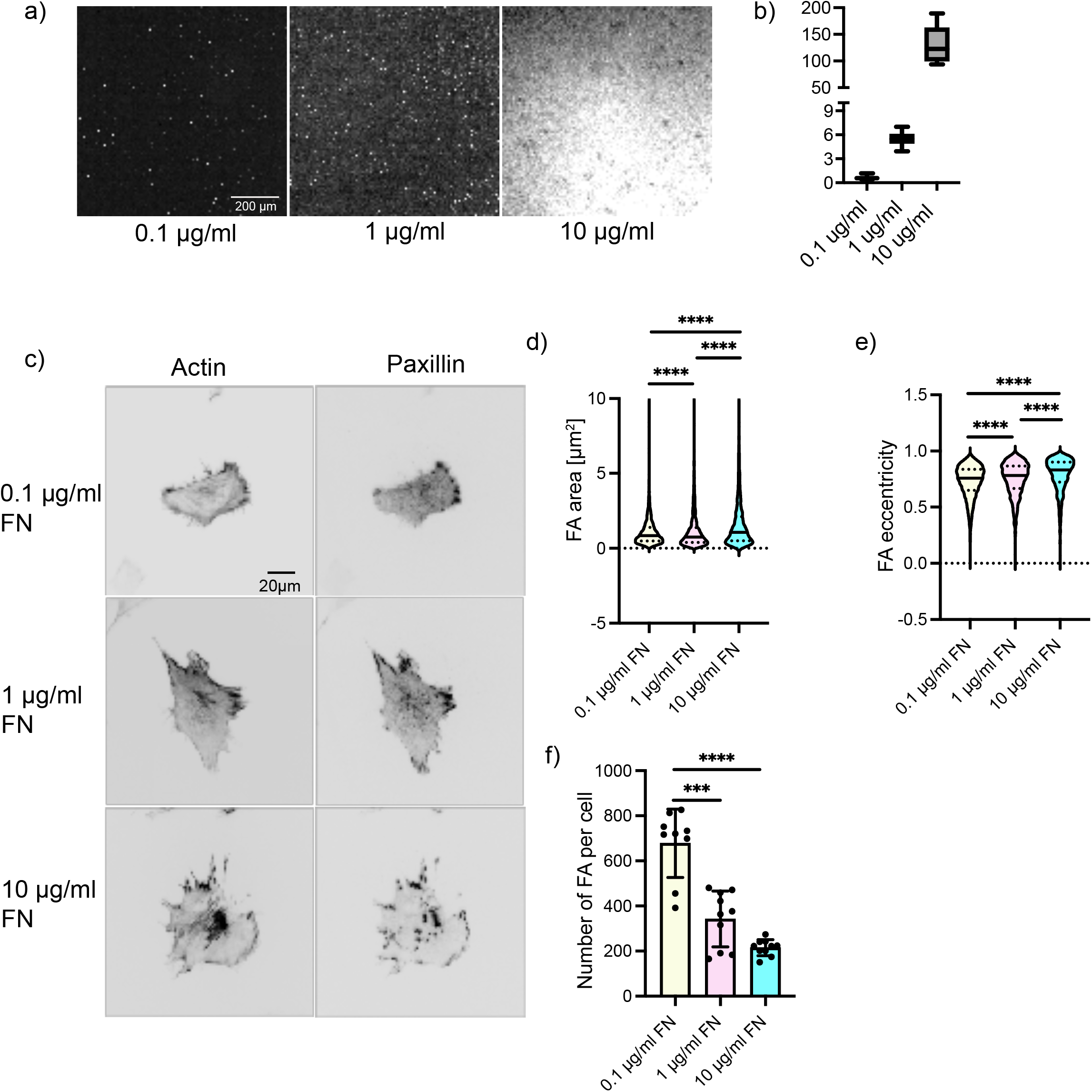
(a) Representative images of glass bottom dishes coated with different concentrations of fibronectin (0.1 ug/ml, 1ug/ml, 10ug/ml) (b) box plot of mean fibronectin intensities at the three different concentrations. (c) Representative images of MEFs plated on glass coated with different FN concentrations (0.1 ug/ml, 1ug/ml, 10ug/ml), fixed, and stained with phalloidin 488 to label actin and Alexa 568 to label paxillin. Violin plots of FA area (d) FA eccentricity (e) and box plot of number of FA per cell (f) in MEFs plated on glass coated with different FN concentrations (0.1 ug/ml, 1ug/ml, 10ug/ml). Quantifications were from analysis of immunofluorescence images of paxillin. N = 10 for each condition.

**Figure S2.**
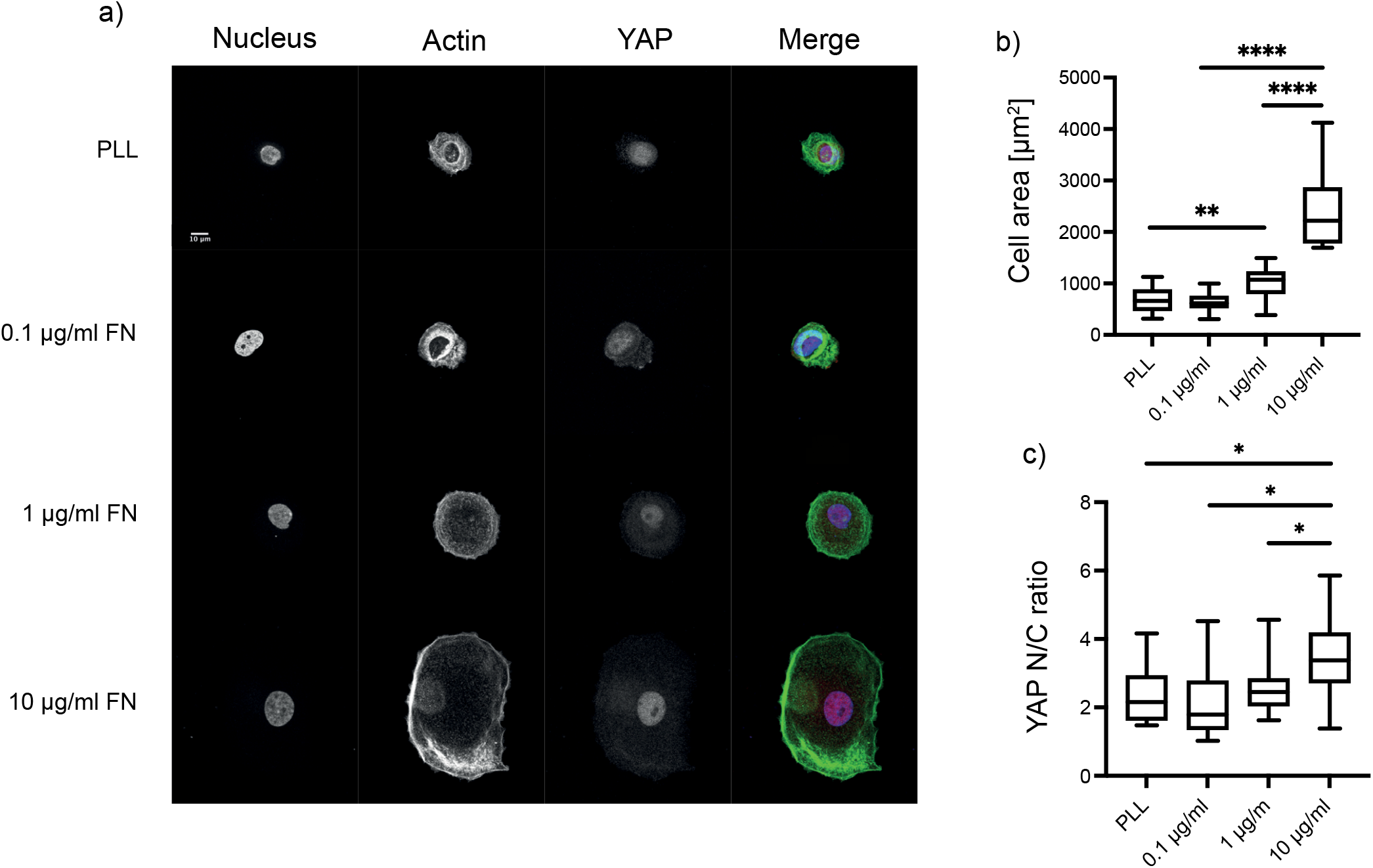
(a) Representative images of MCF10a cells plated on glass coated with different FN concentrations (0.1 ug/ml, 1ug/ml, 10ug/ml), fixed and stained with Hoechst to stain nucleus, phalloidin 488 to label actin and Alexa 568 to label YAP. (b) Box plot quantification of cell area of MCF10a cells from analysis of immunofluorescence images of cells plated on glass coated with different FN concentrations (0.1 ug/ml, 1ug/ml, 10ug/ml) and PLL. Cell area was obtained by segmenting the actin channel. N = 13 for each condition. (c) Box plot quantification of nuclear/cytoplasmic ratio of YAP of MCF10a cells from analysis of immunofluorescence images of cells plated on glass coated with different FN concentrations (0.1 ug/ml, 1ug/ml, 10ug/ml) and PLL. N = 13 for each condition.

**Figure S3.**
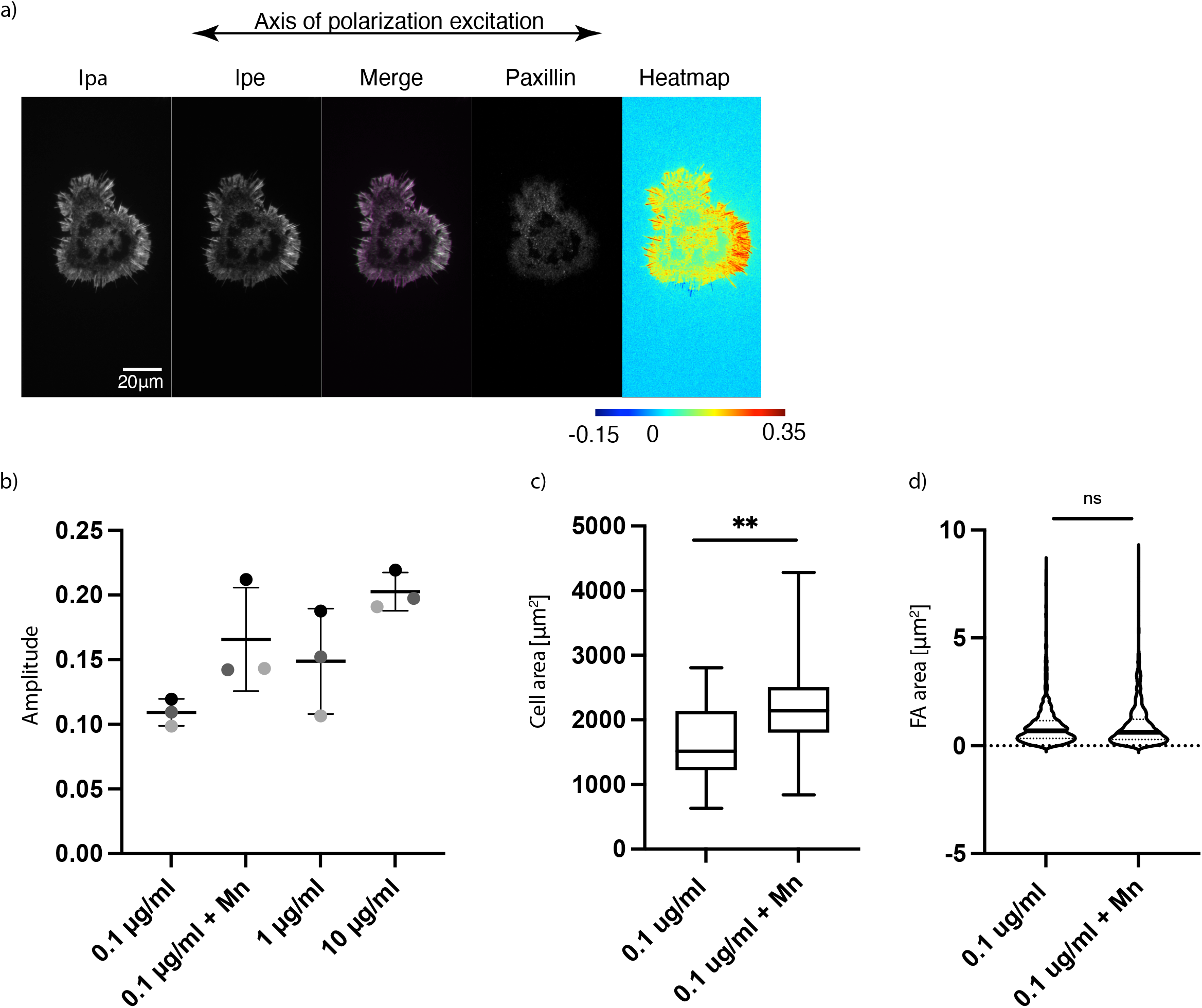
(a) Representative images of MEFs plated on glass coated with PLL. Immunofluorescence images of phalloidin 488 actin with emission from the parallel (Ipa) and perpendicular (Ipe) channels and merge are shown. Paxillin stained with Alexa 568 (middle). Emission anisotropy (r) of actin stained with phalloidin 488 in the whole cell is shown (right). (b) Graph showing amplitudes of actin anisotropy (r) in cells plated on glass coated with different FN concentrations (0.1 ug/ml, 1ug/ml, 10ug/ml) and in the presence of Mn^2+^ on 0.1 ug/ml of FN from 3 experiments. (c) Box plot quantification of cell area from analysis of immunofluorescence images of cells plated on glass coated with 0.1 ug/ml of FN and in presence or absence of Mn^2+^. Cell area was obtained by segmenting the actin channel. N = 36 for each condition. (d) Violin plot quantification of nuclear/cytoplasmic ratio of YAP from analysis of immunofluorescence images of cells plated on glass coated with 0.1 ug/ml of FN and in presence or absence of Mn^2+^. N = 12 for each condition.

**Figure S4.**
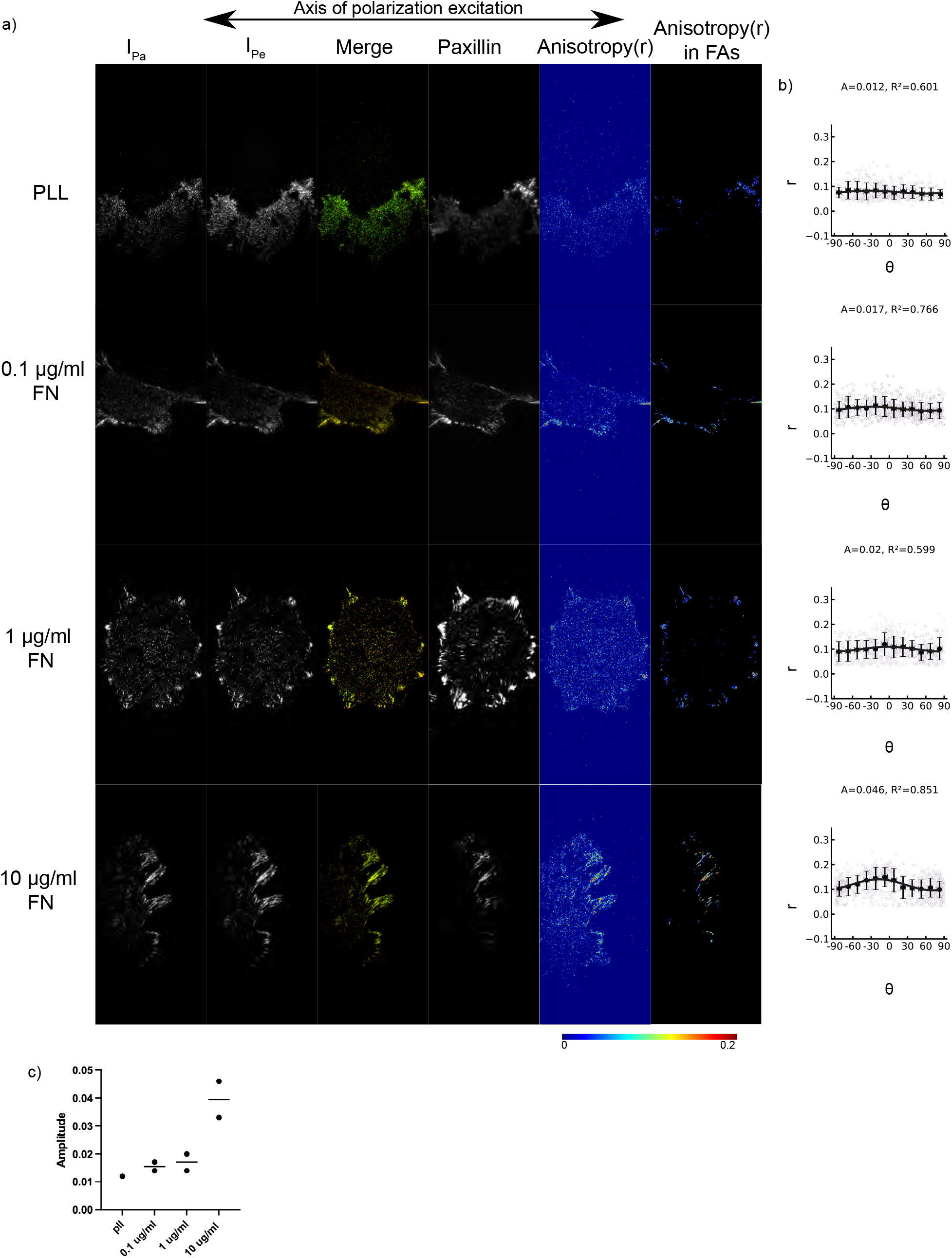
(a) Representative images of aV-integrin-GFP transfected MEFs plated on glass coated with PLL and different FN concentrations (0.1ug/ml, 1ug/ml 10ug/ml) imaged with TIRF. Emission from the parallel (Ipa) and perpendicular (Ipe) channels and merge are shown (left Ipa magenta, Ipe green). Paxillin stained with Alexa 568 (middle). Emission anisotropy (r) of actin stained with aV-integrin-GFP in the whole cell and in the FAs are shown (right). Magnitude of anisotropy color scale (bottom) (b) mean integrin anisotropy (r) in FAs vs FA orientation fit to the cos2 function r = C + A-cos2(y + 8d) for cells plated on each PLL/FN condition. Error bars represent SD. N = 15 for each experiment. (c) graph of integrin anisotropy amplitudes of cells on different FN concentrations (0.1ug/ml, 1ug/ml 10ug/ml) from two experiments. PLL experiments were done once. N = 19, 80, 35, 26 for the conditions PLL, 0.1ug/ml FN, 1ug/mlFN, 10ug/ml FN respectively.

**Figure S5.**
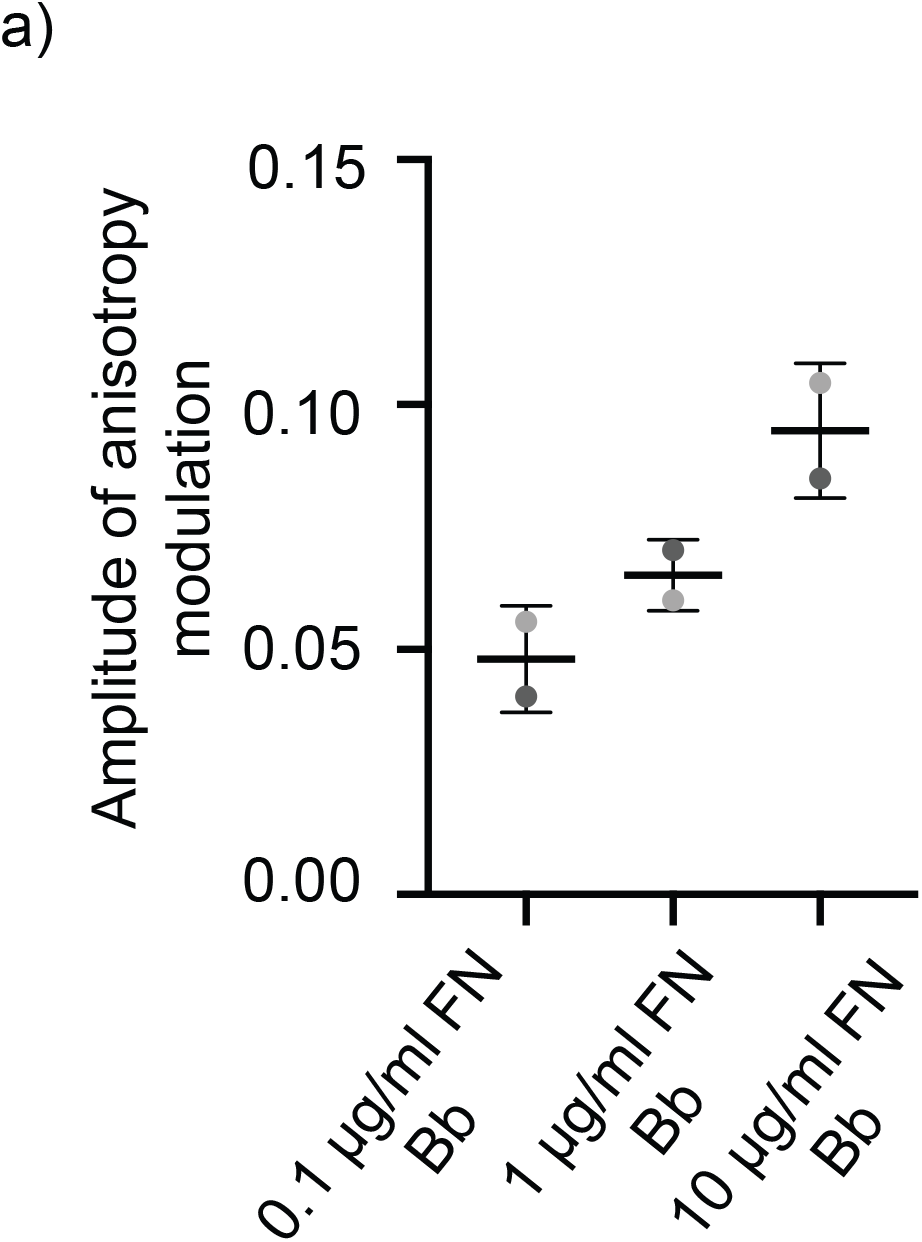
(a) Graph showing amplitudes of actin anisotropy (r) of cells plated on glass coated with different FN concentrations (0.1 ug/ml, 1ug/ml, 10ug/ml) following blebbistatin washout. N = 15 for each condition per experiment.

**Figure S6.**
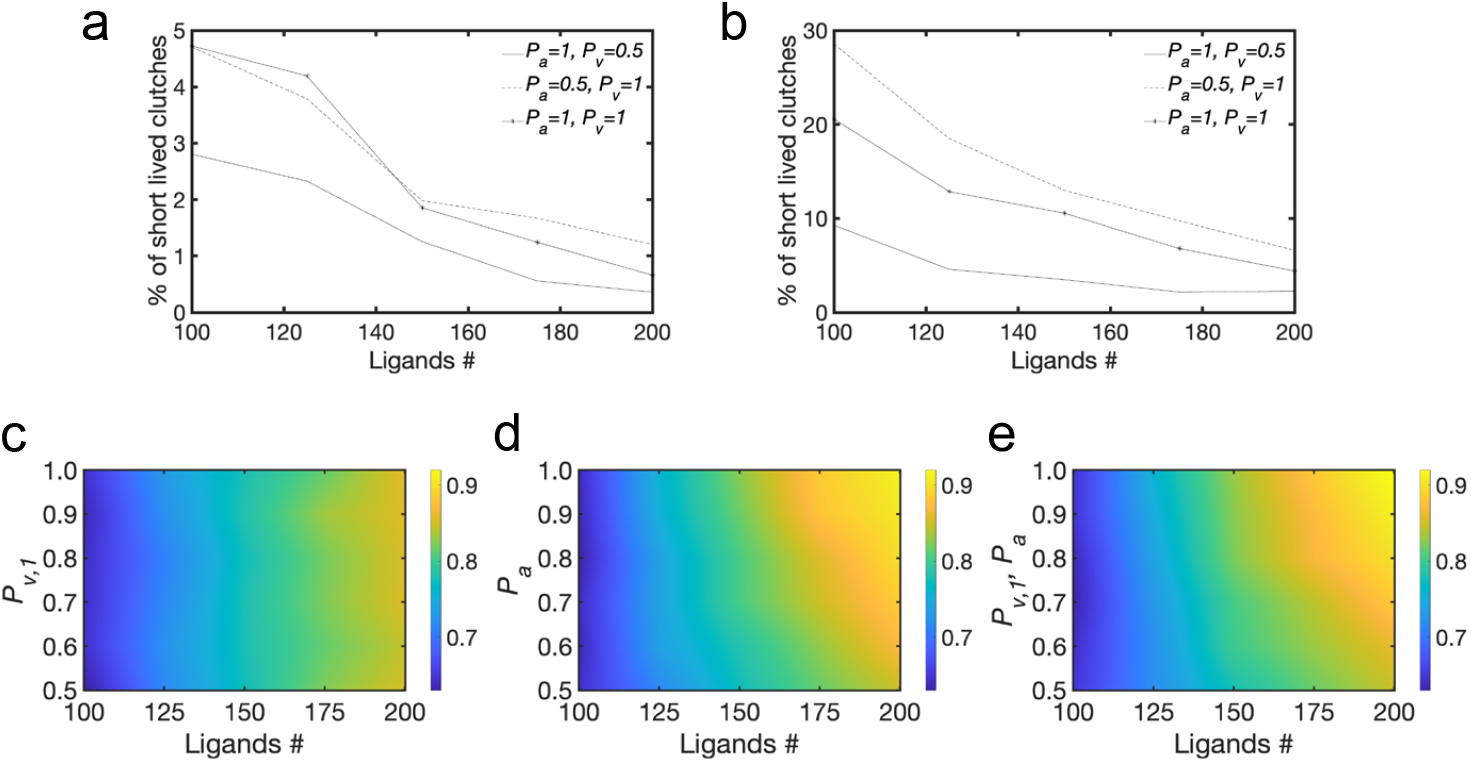
(a) Average percentage of short lived clutches (with lifetime < 0.9 s) relative to the total amount of ligated clutches for different conditions of *P*_*a*_ and *P*_*v,1*_ using 100 to 200 ligands/um^2^ and *Y* = 6 KPa or (b) *Y* = 1 KPa. (c-e) Average fraction of ligated clutches, using *Y* = 1 KPa and varying *n* between 100-200 ligands/um^2^ and either (c) varying *P*_*a*_ between 0.5-1 while keeping *P*_*v,1*_ = 0.5; (d) varying *P*_*a*_ between 0.5-1 while keeping *P*_*v,1*_ = 0.5; (e) simultaneously varying *P*_*v,1*_ and *P*_*a*_ between 0.5-1. All data are obtained as averages from 300 s of simulations for each condition

## Supplementary Note on Computational model

We used a Brownian Dynamics approach to simulate the binding and unbinding of molecular adhesion clutches in response to differences in actin filaments orientations, ligand density and substrate stiffness. Actin filaments were considered implicitly in the model, by modulating the amount of force they exerted on the ligand-bound clutches depending on the number of engaged clutches and molecular motors. Therefore, force on the clutches depended on the actin flow rate, which in turn depended on the number of ligand-bound clutches, substrate rigidity and the number of motors. Unbinding of the clutches could follow two pathways: the direction and the non-directional bond kinetics of vinculin. To simulate different degrees of orientation of the actin filaments, we modulated the probability of establishing a directional versus a non-directional catch bond for the interaction of the clutch with actin.

### Computational Domain and Boundary Conditions

Like our previous implementation of the adhesion assembly model[1], [2], the computational domain was 3D and consisted of two parallel surfaces of 1 μm side, separated in the vertical direction by *L*=20 nm, a dimension typical of integrin headpiece extension[3]. The bottom surface represented the substrate with immobilized ligands; the top substrate represented the ventral cell membrane with the clutches on it (Figure 5b). The clutches were initially randomly distributed on the top surface, and they diffused along *x* and *y*, with diffusion coefficient of integrin, *D* = 0.29 μm^2^/s[4]. To avoid finite size effects on the motion of the free clutches, we used periodic boundary conditions in the lateral directions of the domain.

### Substrate Model

The substrate was considered an elastic solid, consisting of a randomly distributed ideal linear springs, mimicking individual fibronectin molecules, with stiffness depending on substrate rigidity, as:

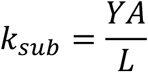

where *Y* is the Young’s modulus (we tested values in the range 0.4– 60 kPa), A is the integrin/ligand cross-sectional area (corresponding to 80 nm^2^, from an ideal bar of radius ∼5 nm, corresponding to approximately half the value of the transmembrane leg separation of an integrin in the open conformation), and *L* is the equilibrium distance separation between substrate and top layer, corresponding to the equilibrium separation between clutch and bound ligand.

Hooke’s law for each spring in the bundle can be written as:

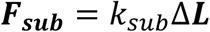

where Δ***L*** is the variation from the equilibrium separation between clutch and bound ligand. We used ligand densities between 100-200 ligands/μm^2^.

### Actin flow

Actin was considered implicitly, as a force acting on each ligated clutch, as a function of the flow velocity. It was calculated as:

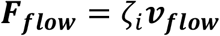

The actin retrograde flow velocity (***v***_***flow***_) was calculated through the linear force-velocity relationship[5], [6],:

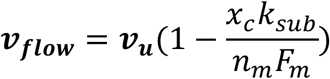

where ***v***_***u***_ = 0.11 μm /s is the unloaded velocity, *x*_*c*_ is the number of bound clutches, *n*_*m*_ =135 is the number of motors, and *F*_*m*_ =2 pN is the motor stall force.

### Clutch Representation

Each *i*-th clutch and *j*-th ligand was defined by a 3D position vectors, ***r***_***i***_ and ***r***_***j***_, respectively. The vector ***r***_***i***_ presented *x, y*, and *z* coordinates of the *i*-th clutch; the vector ***r***_***j***_ presented *x, y* and *z* coordinates of the *j*-th ligand. At every timestep of the simulations, *x* and *y* of ***r***_***i***_ were updated to track the clutch displacement on the top surface, and binding/unbinding of substrate ligands, while *z* remained fixed at 0; *x, y*, and *z* of ***r***_***j***_ remained all fixed over the course of the simulations because ligands were immobilized, with *z* = -0.02 μm.

When a free, diffusive clutch came in proximity of a free ligand (< 21 nm from it), it bound the ligand by establishing a harmonic interaction. This interaction presented spring constant *k*_*sub*_ (proportional to the substrate Youngs’ modulus, *Y*), and equilibrium distance *L*, corresponding to the separation of the open extracellular integrin headpiece from the membrane [3]. In the ligated state, a number of vinculin-actin bonds were also formed. This number dependent on the force on the integrin-fibronectin bond: 2 for forces below 8pN; 5 for forces between 8 and 15 pN; 9 for forces between 15 and 21 pN; 11 for forces > 21 pN [7]. The harmonic interactions between clutches and ligands, and between vinculins and actin provided several mechanical links between actin and the substrate. Each of these links lasted for a certain lifetime, that depended on two types of catch-slip bonds (integrin-ligand and vinculin-actin bonds). Each clutch became unligated if integrin and all vinculins failed. Once the clutches unbound their ligands, they became diffusive again until they bound a new ligand.

### Implementation Algorithm

Recognizing that inertia is negligible on the length and time scales of integrin motion in the plasma membrane, the displacement of each *i-*th integrin was governed by the Langevin equation of motion in the limit of high friction[8]

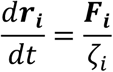

where ***r***_***i***_ was the position vector of integrin; *ζ*_*i*_ was integrin friction coefficient calculated using Einstein relation, as 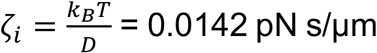, where *k*_*B*_*T* = 4.11 pN nm and *D* = 0.29 μm^2^/s; *dt* = 10^−4^ s was the simulation timestep; *F*_*i*_ was the total force on the clutch, including a stochastic contribution from thermal effects and a deterministic contribution from actin flow, governed by amount of ligated clutches, number of motors and their stall force, and substrate mechanics.

Considering all forces acting on the clutches at every time step, their positions were updated iteratively using an explicit Euler integration scheme:

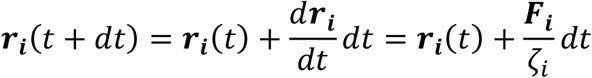

### Forces acting on the clutches

The total force acting on each *i*-th clutch, ***F***_***i***_, resulted from the sum of a stochastic and a deterministic contribution. A stochastic force, ***F***_***T***_, was applied to all clutches in the membrane at every *dt*, to mimic thermal effects generating diffusion. This force satisfied the fluctuation-dissipation theorem. Deterministic forces originated from actin flow, ***F***_***flow***_, and substrate tension, ***F***_***sub***_. Thus, the total force acting on each *i*-th clutch was calculated as:

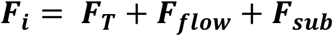

### Lifetime of clutch engagements depends on force

The clutch/substrate interaction persisted for a characteristic lifetime which depended upon the tension on the bond. We assumed that integrin and vinculin feel the effect of this force equally on the clutch bond and that the force increases the dissociation rate constants (k_off,integrin_, k_off,+_, k_off,-_) as originally proposed by Bell [9].

Two characteristic catch bonds were considered: integrin-fibronectin and vinculin-actin. The clutch/substrate interaction broke when both bonds were broken (for the case of vinculin, all vinculin bonds had to break for the clutch to become unligated).

A double exponential pathway determined unbinding rates of integrin-fibronectin and of each vinculin-actin. It included a strengthening pathway, with a negative exponent, and a weakening pathway, with a positive exponent.

For integrin, the unbinding rate was:

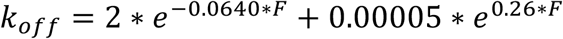

For each vinculin, the unbinding rate depended on filaments orientation and therefore it depended on the direction of force application. For forces applied toward the filament pointed end, the catch bond was directional, with the longest lifetimes and corresponding unbinding rate:

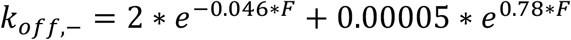

For forces applied toward the filament barbed end, the catch bond was non-directional, and the unbinding rate was:

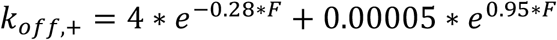

